# Fentanyl and fentanyl-xylazine co-exposure influence mitochondrial transcriptional programs in human trophoblasts

**DOI:** 10.64898/2026.10.31.763373

**Authors:** Jiayu Meng, Lei Liu, Sheng Liu, Keyi Huang, Yoel Sadovsky, Yingshi Ouyang

## Abstract

**Introduction:** Illicit fentanyl exposure during pregnancy is associated with adverse outcomes. Yet its direct effects and their underlying molecular mechanisms in human trophoblasts remain largely unknown. Xylazine and medetomidine, two ɑ2-adrenergic agonists, were recently detected as adulterants in illicit fentanyl-containing products, and have been linked to worsened clinical outcomes.

**Methods:** We assessed the transcriptional responses of human trophoblast stem (TS) cells in their cytotrophoblast state (TS-CTBs) and derived syncytiotrophoblasts (STBs) after 4-day and 9-day exposure to fentanyl alone or combined with medetomidine and xylazine (FMX).

**Results:** Fentanyl alone produced modest gene expression changes in TS-CTBs, but consistently induced suppression of mitochondrial and ribosomal transcriptional programs in STBs regardless of exposure duration, an effect absent in parental TS-CTBs. 9-day FMX co-exposure influenced mitochondrial respiratory chain gene expression in derived STBs, resulting in discordant regulation of nuclear *versus* mitochondrial genome-encoded respiratory chain genes. Temporal pathway analysis reveals progressive trophoblast remodeling and dysregulation of mitochondrial transcriptional programs following fentanyl and FMX co-exposure.

**Discussion:** Together, these findings illuminate trophoblast pathways affected by fentanyl alone and by FMX co-exposure, and establish a basis for future studies examining the functional and clinical relevance of these transcriptional responses.

**Highlights:**

- Syncytiotrophoblasts are transcriptionally sensitive to fentanyl exposure
- Mitochondrial transcriptional programs are suppressed in drug-exposed trophoblasts
- FMX co-exposure reshapes fentanyl-associated transcriptional changes in trophoblasts

## INTRODUCTION

The overall prevalence of exposure to opioids during pregnancy, and specifically to fentanyl, has markedly increased in recent decades, raising major public health concerns^1^. Prenatal opioid exposure has been associated with adverse pregnancy outcomes, including substandard fetal growth, preterm delivery and stillbirth^2–4^. Infants exposed *in utero* are at increased risk of neonatal opioid withdrawal syndrome and may exhibit neurodevelopmental impairments in the long term^5–7^. Although these outcomes are often attributed to maternal factors, the impact of these drugs on placental cellular responses remains poorly defined. Alarmingly, illicit fentanyl has been increasingly adulterated with xylazine and medetomidine, two α2-adrenergic agonists commonly used in veterinary medicine^8–10^. There have been several reports of gestational exposure to these agents^11,12^, associated with skin injury, as well as with prolonged, severe withdrawal syndrome^13,14^. However, the effect of these combined exposures on placental biology at the molecular level remains unknown.

Previous studies investigating opioid exposure during pregnancy have largely relied on placental RNA-seq^15,16^, providing limited resolution regarding cell-type–specific responses within the placenta. Given the central role of villous trophoblasts in placental function, understanding how these cells transcriptionally respond to opioid and non-opioid sedative agents represents a critical knowledge gap.

Here, we used human trophoblast stem cells derived from first-trimester cytotrophoblasts^17^ to determine whether the addition of xylazine and medetomidine modifies the cell-state- and time-dependent transcriptional responses to fentanyl exposure. Transcriptomes were analyzed in cells maintained in a stem/progenitor state (TS-CTBs) or differentiated into syncytiotrophoblasts (TS-STBs), following 4-day or 9-day exposure to fentanyl alone or to fentanyl combined with medetomidine and xylazine. We identified shared and distinct transcriptional responses across trophoblast states, exposure durations, and exposure conditions. To our knowledge,this is the first transcriptome-wide study of villous trophoblast responses to fentanyl combined with xylazine and medetomidine, providing a molecular framework for investigating the placental cellular consequences of this emerging polysubstance-exposure pattern.

## METHODS

### Cell Culture

Human TS cells CT30 clone derived from first-trimester placentas were cultured as previously described and obtained from the RIKEN Cell Bank^17^. Culture dishes were pre-coated with 5 ug/ml Collagen IV (Sigma, #C5533-5MG) diluted in PBS (ThermoFisher, #14190250) supplemented with 0.5% penicillin–streptomycin (ThermoFisher, #15140122) for at least 1 h at 37°C in 5% CO_2_.

To maintain TS cells in an undifferentiated state, cells were grown in stemness medium consisting of DMEM/F-12, GlutaMAX (ThermoFisher, #10565042) supplemented with 0.5% penicillin–streptomycin, 0.3% bovine serum albumin (BSA; Sigma, #A9576-50ML), 1% ITS-X supplement (ThermoFisher, #51500056), 0.1 mM 2-mercaptoethanol (ThermoFisher, #31350010), 0.2% FBS (ThermoFisher, #16141079), 1.5 μg/ml L-ascorbic acid (Sigma, #A4544- 25G), 50 ng/ml human epidermal growth factor (EGF; Peprotech, #AF-100-15), 1 μM SB431542 (Selleckchem, #S1067, 10 mg), 0.5 μM A83-01 (Sigma, #SML0788-5MG), 0.8 mM Valproic acid (Sigma, #P4543-10G), 2 μM CHIR99021 (Selleckchem, #S1263), 5 μM Y-27632 (Selleckchem, #S1049, 10 mg). TS cells were maintained at 37°C in a humidified incubator with 5% CO_2_ and passaged every 3-4 days at a split ratio of 1:3. For passaging, cells were washed with PBS and dissociated using TrpLE Express (ThermoFisher, #12604013) at 37°C in 5% CO_2_.

For differentiation, cell culture plates were pre-coated with 2.5 ug/ml Collagen IV for at least 1h at 37°C in 5% CO_2_. To induce differentiation into STBs, TS cells were cultured in differentiation medium consisting of DMEM/F-12, GlutaMAX supplemented with 0.5% penicillin–streptomycin 0.3% BSA, 1% ITS-X supplement, 0.1 mM 2-mercaptoethanol, 4% knockout serum replacement (KSR; ThermoFisher, #10828028), 2 μM forskolin (Selleckchem, #S2449), 2.5 μM Y-27632.

Exposure during differentiation was limited to 2 days to examine transcriptional responses during early STB formation (Fig. S1).

### Drug exposure

To assess the effects of fentanyl, medetomidine, and xylazine on trophoblast transcriptomes, TS cells were exposed to 1 μM fentanyl (Hikma, #00641-6027-25) alone or in combination with 100 nM medetomidine (Selleckchem, #S3060) and 1 μM xylazine (Sigma, #1720429) under either stemness or differentiation conditions, as illustrated schematically in Figs.1A, 2A, 3A, and 4A.

Fentanyl citrate injection (Hikma Pharmaceuticals, 50 µg/mL, NDC 0641-6027-25) was added directly to the culture medium to a final concentration of 1 µM. Medetomidine HCl (Selleckchem, #S3060) and xylazine hydrochloride (Sigma, #1720429) was dissolved in sterile nucleas-free water to make 10 µM and 100 µM stock stocks, respectively, and diluted to final concentrations of 100 nM and 1 µM, respectively. For FMX exposure, the three drugs were added simultaneously to the culture medium as separate additions. Vehicle controls received sterile water in volumes matched to each treatment, including the combined volume for FMX. Medetomidine and xylazine stock solutions were stored at −20°C. The commercial fentanyl solution was stored at room temperature according to manufacturer’s instructions. These drug concentrations were applied in both 4-day and 9-day exposure paradigms. Drugs and media were replenished every 24 h.

For 4-day exposure, TS cells were seeded at 3–4 x 10^5^ cells per well in 6-well plates and exposed for 2 days in stemness medium. On day 2, cells were split half. One half was maintained in stemness medium with continued drug exposure for an additional 2 days, while the other half was switched to differentiation medium with drugs for 2 days.

For 9-day exposure, TS cells were seeded at 3–4 x 10^5^ cells per well in 6-well plates and exposed for 7 days in stemness medium. On day 7, cells were split half. One half was maintained in stemness medium with continued drug exposure for an additional 2 days, while the other half was switched to differentiation medium with drugs for 2 days.

Drug concentrations were selected as *in vitro* exposure conditions, which was informed by clinical and forensic toxicology reports of fentanyl and adulterant co-exposure^18–20^. The concentrations were not intended to directly reproduce typical circulating maternal plasma concentrations. The single-concentration design was not intended to define concentration-response relatioships, threshold concentrations, or transcriptomic points of departure.

### RNA extraction, library preparation, and RNAseq statistical analyses

The RNA-seq datasets comprise 42 samples from three independent experimental batches, each with its own vehicle controls and drug-exposed biological replicates (n=3 per condition). Batch 1 included TS-CTBs and STBs exposed for 4 days to vehicle, fentanyl (Fen), or the fentanyl/medetomidine/xylazine combination (FMX). Batch 2 included TS-CTBs and STBs exposed for 9 days to vehicle or Fen. Batch 3 included TS-CTBs and STBs exposed for 9 days to vehicle or FMX. Vehicle controls were matched to each cell state and exposure duration within each batch (Fig. S3-S5). Total RNAs were extracted on day 4 or day 9, using TRIzol RNA isolation reagent (Thermo Fisher, #15596026), followed by purification with the RNeasy mini kit (Qiagen, #74104) including on-column RNase-free DNase (Qiagen, #79254), according to the manufacturer’s protocols as previously described^16^. RNA quality was assessed using an Agilent HS Total RNA 15-nt kit (Agilent, DNF-472T33) on an Agilent Analytical 5300 Fragment Analyzer. Samples with RNA integrity number (RIN) ≥7 were used for library preparation.

RNAseq libraries were prepared by Novogene using poly (A) selection and sequenced on an Illumina NovaSeq X plus platform to generate paired-end 150-bp reads, with an average depth of 40 million individual reads per sample. After quality control, clean reads were aligned to the human reference genome (GRCh38) using STAR^21^ with default parameters. Uniquely mapped sequencing reads were assigned to GENCODE v47^22^ genes using featureCounts^23^ with the parameters “–p –Q 10 –O”. Genes with read counts more than 10 in at least three of the samples were retained for downstream analysis. Count data were normalized using the TMM (trimmed mean of M values) method and differential gene analysis was performed using edgeR (v3.20.8)^24,25^. P values were adjusted using the Benjamini-Hochberg procedure^26^ to control the false discovery rate (FDR), and genes with an adjusted p value less than 0.05 were considered significantly differentially expressed (Fig. S2).

### Gene set enrichment analysis

Gene Set Enrichment Analysis (GSEA) was performed to identify coordinately regulated biological pathways. It was used to determine whether predefined gene sets exhibited coordinated enrichment across the entire ranked gene list, thereby capturing pathway-level changes that may not be detected by DEG-based analyses. Genes were pre-ranked using the metric -log10(FDR) x sign (logFC), where sign(logFC) indicates the direction of change (+1 for upregulated genes and −1 for downregulated genes). This is a widely adopted approach that integrates statistical significance with direction of expression change^27^. The GSEA was executed using the gseGO function from the clusterProfiler R package^28^. Functional enrichment was tested against Gene Ontology (GO) terms^29^, using the human genome annotation (org.Hs.eg.db)^30^. Gene sets with sizes outside the range of 10 to 500 were excluded to ensure biological relevance and statistical robustness. Multiple testing correction was applied internally by *clusterProfiler*, and gene sets with an adjusted p value below 0.1 were considered significant. Analyses were run under a fixed random seed with serialized processing to ensure reproducibility.

### Gene set variation analysis

Gene Set Variation Analysis (GSVA) was performed using the GSVA R package to estimate pathway activity at the individual sample level, enabling the assessment of dynamic pathway activity changes across experimental conditions and developmental states^31^. Unlike the GSEA, which evaluates enrichment with respect to a predefined phenotypic contrast and produced a single enrichment statistic per gene set, GSVA operated in an unsupervised, sample-wise manner, generating a continuous enrichment score for each gene set in each sample. This formulation transformed the gene-by-sample expression matrix into a gene set-by-sample enrichment matrix, enabling flexible downstream statistical modeling that was not constrained to pairwise comparisons.

GSVA enrichment scores were computed using a Gaussian kernel density estimator, and differential pathway enrichment between drug-exposed and vehicle control conditions was assessed using the limma linear modeling framework^32^. A design matrix encoding all combinations of cell type, drug exposure, and time point was constructed, and pairwise contrasts were defined to isolate drug effects within each cell type and time point. Empirical Bayes moderation was applied via eBayes, and pathways with a Benjamini–Hochberg-adjusted p-value less than 0.1 in at least one contrast were considered significant and retained for visualization and temporal pattern analysis.

To identify common patterns of pathway responses across experimental conditions, differential pathway activity was summarized as ΔGSVA score (drug – control). Each pathway was represented by an 8-dimensional vector corresponding to its ΔGSVA scores across the eight exposures. Pathways were grouped according to their response profiles using k-means clustering with k = 8 and a random seed of 42. K-means clustering partitions pathways into clusters by minimizing within-cluster variance. Briefly, pathways were assigned to the nearest cluster centroid based on Euclidean distance, centroids were updated as the mean response profile of all pathways within each cluster, and assignment and update steps were repeated until convergence. The resulting clusters represent distinct patterns of pathway regulation, capturing differences in temporal responses, cell-type specificity, and drug selectivity across the experimental conditions.

### Gene Ontology (GO) Enrichment Analysis

Gene Ontology (GO) enrichment analysis was performed to identify biological processes, molecular functions, and cellular components associated with differentially expressed genes. DEGs were filtered using a false discovery rate (FDR) of 0.05. Upregulated and downregulated DEG sets were separately defined and gene ontology enrichment analysis was performed independently for each DEG set across all three ontologies (Biological Process, Cellular Component, and Molecular Function) using the *enrichGO* function in *clusterProfiler*^28^, with Benjamini-Hochberg multiple testing correction applied^26^ and adjusted p-value cutoff of 0.1.

## RESULTS

### Distinct transcriptional changes of TS-CTBs to 4-day and 9-day fentanyl exposure

To assess the effect of fentanyl exposure on TS-CTBs, we cultured cells in stemness medium with or without fentanyl for 4 days or 9 days (Fen4 or Fen9, respectively; Fig. 1A). Informed by human toxicology reports, we selected the fentanyl concentration to represent a sustained, high-exposure *in vitro* condition rather than a direct simulation of typical human plasma exposure.

**Figure 1.**
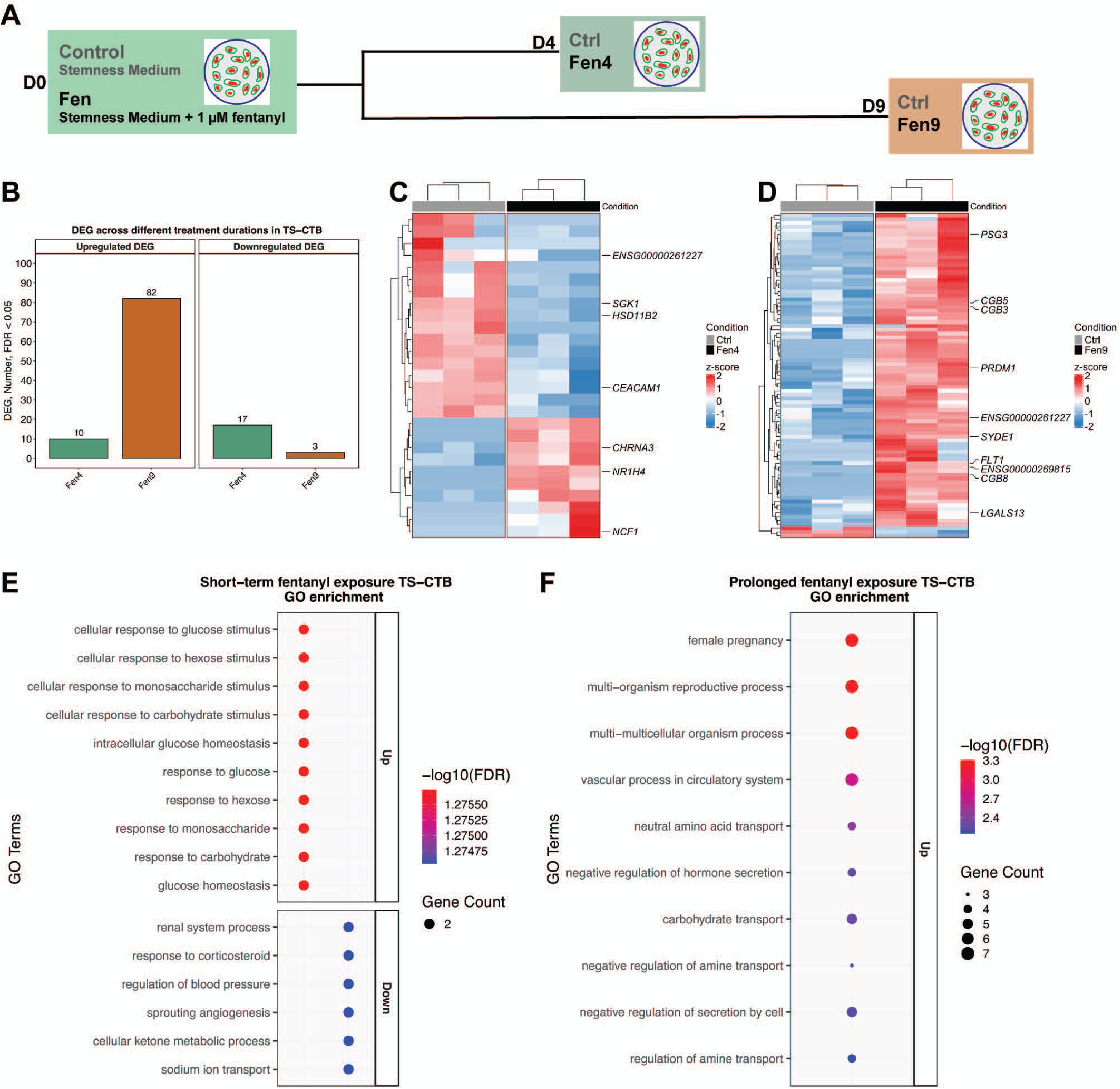
Transcriptional landscape of TS-CTBs under 4-day and 9-day fentanyl exposure. (A) An experimental schematic: trophoblast stem cells, maintained in stemness medium (TS-CTBs), were exposed to 1 µM fentanyl for either 4 days (Fen4) or 9 days (Fen9). (B) Barplot showing numbers of differentially expressed genes (DEGs; FDR < 0.05) in TS-CTBs following 4-day and 9-day fentanyl exposure. (C-D) Heatmaps showing DEGs (FDR < 0.05) in TS-CTBs following (C) 4-day or (D) 9-day fentanyl exposure. (E) GO enrichment of upregulated and downregulated DEGs in TS-CTBs following 4-day fentanyl exposure. (F) GO enrichment results of upregulated DEGs in TS-CTBs following 9-day fentanyl exposure.

Transcriptomic profiling revealed relatively limited transcriptomic changes after 4-day exposure. We found 27 differentially expressed genes (DEGs), including 10 upregulated and 17 downregulated mRNAs (Fig. 1B, Table S1). In contrast, extending exposure to 9 days broadened the transcriptional response, yielding 82 upregulated and 3 downregulated genes in TS-CTBs (Fig. 1B, Table S2). Comparison of DEG sets between the two exposure durations revealed minimal overlap. Only one long non-coding RNA (lncRNA) *ENSG00000261227* of unknown function was shared between Fen4 and Fen9. This lncRNA was downregulated (log_2_FC: -2.81) after 4 days but upregulated (log_2_FC: 2.67) after 9 days of exposure. Collectively, these findings suggest that the transcriptional response to fentanyl differed between exposure durations, with largely distinct sets of genes altered at 4 and 9 days relative to time-matched vehicle controls.

4-day fentanyl exposure was associated with induction of genes related to stress responses and metabolic regulation (Fig. 1C). Among the genes upregulated following Fen4, *NCF1* was strongly induced (log_2_FC: 6.15). NCF1 encodes a core component of the NADPH oxidase complex that drives reactive oxygen species production^33^. Additionally, the expression of *NR1H4* (log_2_FC: 5.77), encoding the transcription factor nuclear farnesoid X receptor that controls metabolic homeostasis^34^, and *CHRNA3* (log_2_FC: 4.70), which encodes neuronal nicotinic acetylcholine receptor^35^, were markedly induced. These findings indicate that fentanyl exposure led to activation of inflammatory-redox signaling and metabolic regulatory pathways.

Conversely, several genes implicated in trophoblast function were downregulated after Fen4 (Fig. 1C). The expression of *HSD11B2*, an enzyme that protects the fetus from excessive exposure to maternal cortisol, was decreased (log_2_FC: -0.73). Additional downregulated genes included *SGK1* (log_2_FC: -0.44) and *CEACAM1* (log_2_FC: -0.77), which both are involved in trophoblast invasion and cell adhesion^36,37^.

9-day fentanyl exposure induced a distinct transcriptional signature enriched for genes associated with placental endocrine function (Fig. 1D). These include *PSG3*^38,39^ (log_2_FC: 1.82), *LGALS13*^40^ (log_2_FC: 0.62), and the hormone chorionic gonadotropin genes *CGB3* (log_2_FC: 0.84), CGB5 (log_2_FC: 0.87), and *CGB8*^41,42^ (log_2_FC: 0.78). We also identified increased expression of *SYDE1* (log_2_FC:1.12) and *PRDM1* (log_2_FC: 1.92). SYDE1, a GCM1 target gene, controls cytoskeleton remodeling and trophoblast invasion^43^. Notably, PRDM1 upregulation is associated with preeclampsia through deregulation of trophoblast invasion and apoptosis^44^.

Gene ontology analysis highlighted a temporal divergence in the transcriptional programs affected by fentanyl. Fen4 exposure in TS-CTBs led to enriched pathways related to glucose metabolism and homeostasis, while suppressing corticosteroid response and sodium ion transport (Fig. 1E, Table. S3, S4). In contrast, Fen9 in TS-CTBs prominently upregulated pathways of hCG production, amino acid and carbohydrate transport (Fig. 1F, Table. S5).

Together, these findings suggest that 4-day fentanyl exposure primarily triggers stress-responsive and metabolic reprogramming in TS-CTBs, whereas 9-day exposure shifts the transcriptional program toward a hormone-producing, pregnancy-associated state.

### FMX co-exposure modifies fentanyl-induced transcriptional changes in TS-CTBs

Illicit fentanyl exposure occurs frequently with xylazine and medetomidine. We therefore examined how the FMX mixture modifies fentanyl-associated transcriptional responses (Fig. 2A). Whereas 4 days exposure to FMX (FMX4) induced 17 upregulated and 28 downregulated transcripts (Fig. 2B, Table S6), extended exposure (FMX9) led to 32 genes upregulated and 34 genes downregulated (Fig. 2B, Table S7).

**Figure 2.**
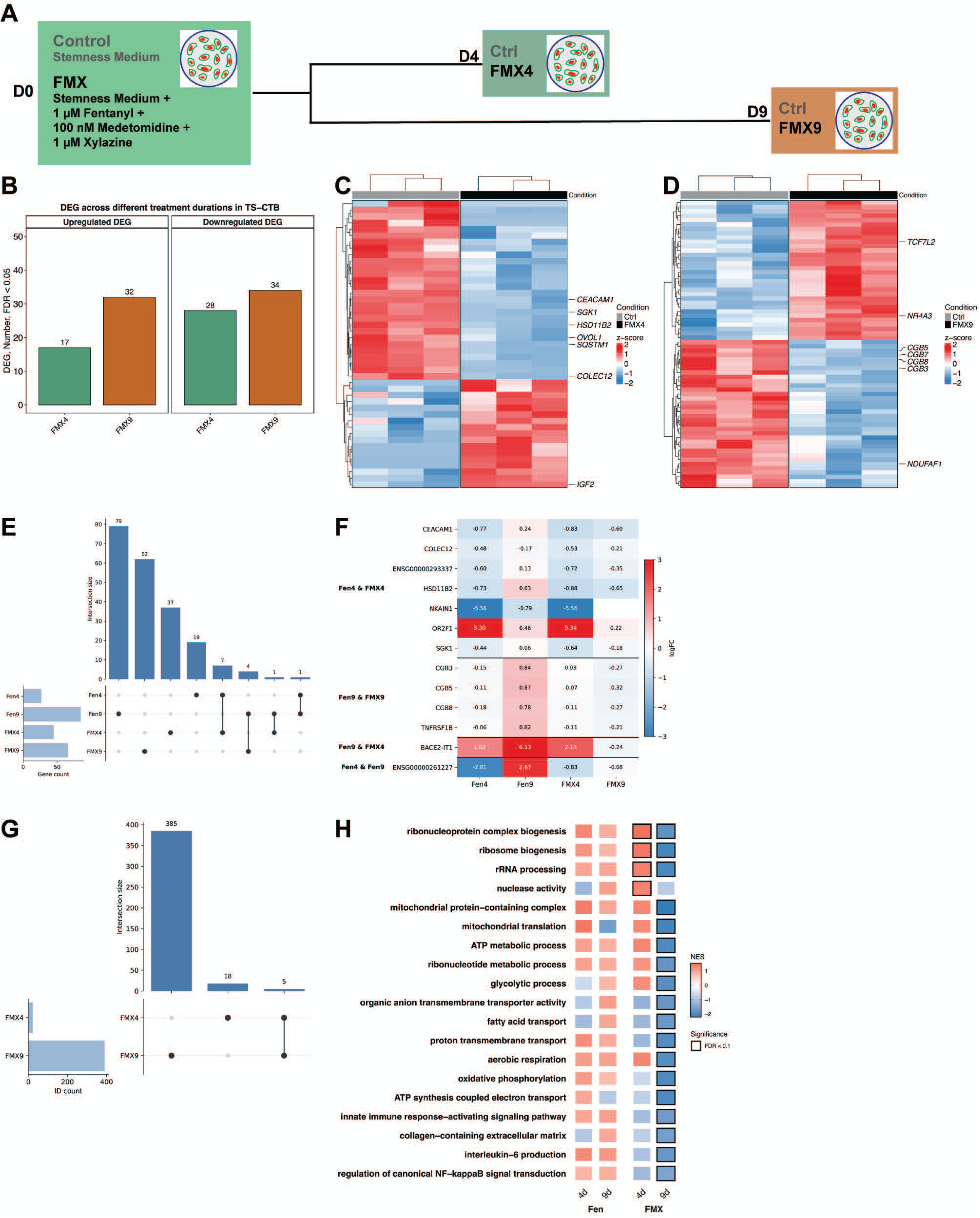
Transcriptional landscape of TS-CTBs under 4-day and 9-day exposure to combined fentanyl-medetomidine-xylazine (FMX). (A) An experimental schematic: TS-CTBs were exposed to 1 µM fentanyl, 100 nM medetomidine, and 1 µM xylazine, for 4 days (FMX4) or 9 days (FMX9). (B) Barplot showing numbers of DEGs (FDR < 0.05) in TS-CTBs following 4-day and 9-day exposure of FMX. (C-D) Heatmaps showing DEGs in TS-CTBs following (C) 4-day and (D) 9-day FMX exposure. (E) Upset plot showing overlapping DEGs between Fentanyl-and FMX-exposed TS-CTBs. (F) Heatmap showing fold-change values of the overlapping DEGs identified in (E). (G) Upset plot showing overlapping enriched pathways between 4-day and 9-day FMX-exposed TS-CTBs. (H) Multi-group comparison of GSEA results. The significantly enriched pathways (FDR < 0.1) were highlighted.

After 4-day exposure to TS-CTBs, the Fen and FMX groups shared only one upregulated and six downregulated genes (Fig. 2C). Specifically, IGF2, a key driver of placental and fetal growth^45^, was upregulated. The downregulated genes included *HSD11B2*, *SGK1*, *CEACAM1*, *COLEC12, OVOL1,* and *SQSTM1*. Note that *COLEC12* is highly expressed in CTBs that mediate innate immune responses^46,47^. *OVOL1* is required for trophoblast differentiation^48^.

Following 9-day exposure to FMX, TS-CTBs exhibited additional transcriptional changes that might influence trophoblast differentiation and mitochondrial metabolism (Fig. 2D). The upregulated genes included *TCF7L2*, a primary effector of Wnt pathway essential for extravillous trophoblast differentiation and invasion^49^, and *NR4A3*, which inhibits trophoblast syncytialization^50^. Conversely, downregulated genes included multiple chorionic gonadotropin β subunits (*CGB3*, *5*, *7*, *8*) and *NDUFAF1*, required for mitochondrial complex I formation^51^.

To examine how the FMX mixture modifies fentanyl-induced transcriptional responses in TS-CTBs, we analyzed the overlap of these DEGs between Fen and FMX groups. Overall, FMX co-exposure did not exert a uniform effect across time but instead produced a temporal shift from early transcriptional amplification to later transcriptional reprogramming. 4-day exposure to fentanyl alone (Fen4) and FMX (FMX4) resulted in seven shared DEGs (Fig. 2E), all of which exhibited larger effect sizes in the FMX4 group (Fig. 2F), suggesting that co-exposure amplifies early transcriptional responses to fentanyl exposure. In contrast, only four DEGs overlapped between 9-day fentanyl (Fen9) and FMX (FMX9) exposure (Fig. 2E), exhibiting opposite direction of gene expression changes between Fen9 and FMX9 (Fig. 2F). These results suggest that extended FMX co-exposure did not simply intensify fentanyl-induced effects, but rather reshaped the transcriptional response toward a distinct molecular program. Only one gene, BACE2-IT1, was commonly upregulated between Fen9 and FMX4 groups (Fig. 2F). BACE2-IT1 is a lncRNA within an intron of the BACE2 gene that governs glucose homeostasis^52^.

We further assessed time-dependent pathway responses to the FMX mixture using gene sets enrichment analysis (GSEA). FMX co-exposure led to distinct transcriptional programs at early and late time points, with 9-day exposure producing substantially broader pathway perturbations than 4-day exposure. Consistent with this observation, 4-day and 9-day FMX exposure resulted in 23 and 390 significantly enriched pathways, respectively, with only five shared between the two conditions (Fig. 2G; Table. S8, S9). 4-day FMX co-exposure in TS-CTBs was characterized by positive enrichment of pathways involved in ribosome biogenesis and rRNA processing. In contrast, 9-day exposure reversed these responses and was notably associated with suppression of mitochondrial translation and ATP synthesis (Fig. 2H, right panel). In addition, 9-day FMX co-exposure led to suppression of immune-related pathways in TS-CTBs, including IL-6 production and NF-κB signaling. In contrast, many of these pathways were not affected by fentanyl alone (Fig. 2H, left panel).

### Fentanyl exposure suppresses mitochondrial and ribosomal transcriptional programs in STBs

We investigated the transcriptional responses to fentanyl in differentiated TS-STBs (Fig. 3A). Increasing the duration of fentanyl exposure led to increased numbers of DEGs in TS-STBs (Fig. 3B, Table S10, S11). 4-day exposure to fentanyl in TS-STBs leads to dysregulation of stress-responsive and metabolism-associated genes (Fig. 3C). These include upregulation of *PHLDA1* (log_2_FC: 0.65), a stress-responsive gene involved in apoptosis. It also led to altered expression of two genes related to protein glycosylation and steroid hormone regulation, *SRD5A3* (log_2_FC: 6.09) and *HSD17B11* (log_2_FC: -0.56, Table. S10). Additionally, two genes *CROT* and *TNFRSF19* were downregulated (Table. S10). CROT is critical for mitochondrial function^53^, whereas TNFRSF19 promotes JNK activation^54^.

**Figure 3.**
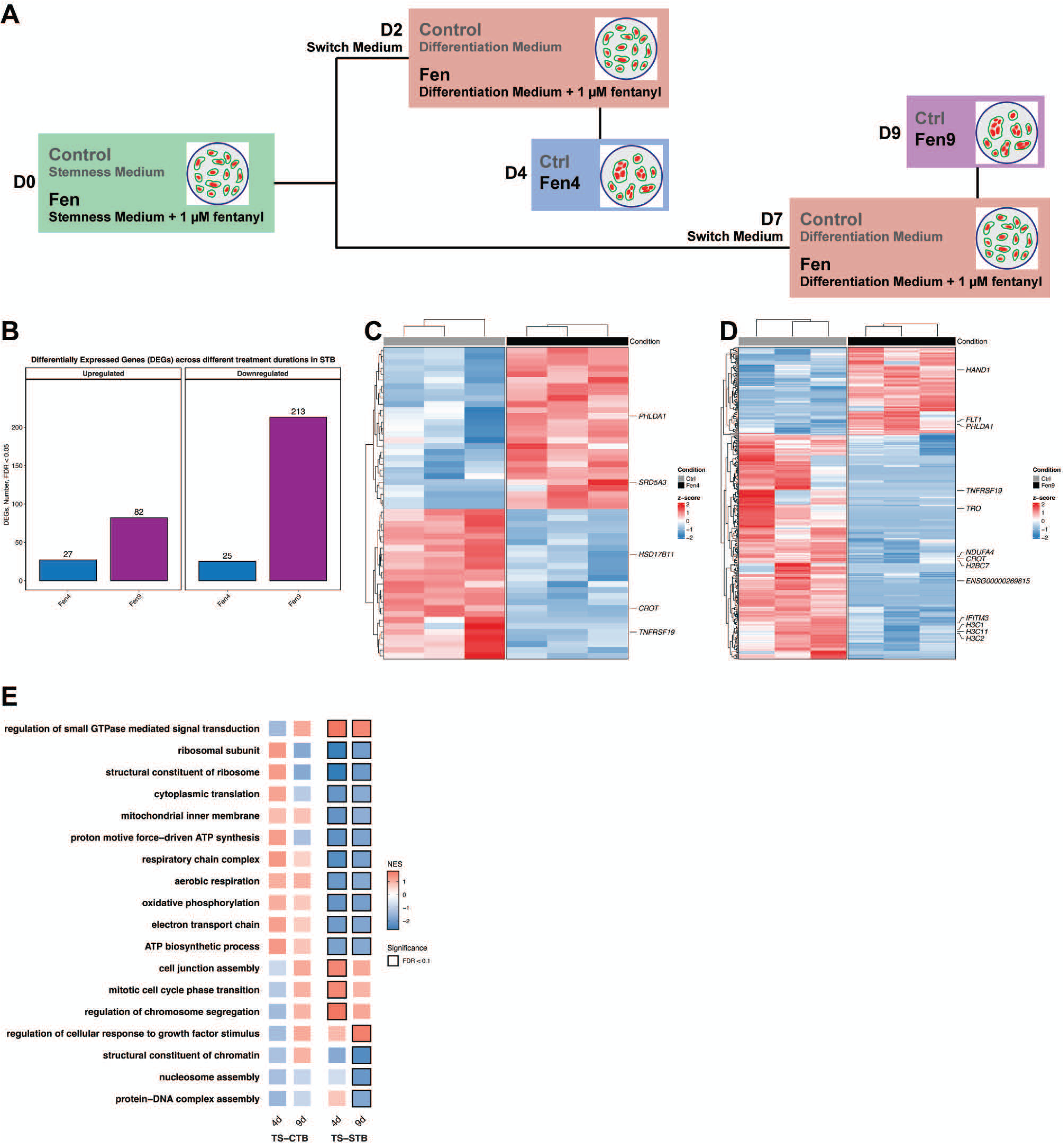
Temporal dynamics of the syncytiotrophoblast (STB) transcriptome under 4-day and 9-day fentanyl exposure. (A) An experimental schematic: TS-CTBs were pre-exposed to fentanyl in stemness medium (2 or 7 days) and subsequently transitioned to fentanyl-containing differentiation medium for 2 days to generate STBs representing 4-day (Fen4) and 9-day (Fen9) total exposure. (B) Barplot showing numbers of DEGs (FDR < 0.05) in 4-day and 9-day fentanyl exposed STB. (C–D) Heatmaps showing DEGs (FDR < 0.05) in STBs differentiated from TS-CTBs, following (C) 4-day and (D) 9-day fentanyl exposure. (E) Multi-group comparison of GSEA results in fentanyl-exposed TS-CTBs and derived STBs at 4 and 9 days. The significantly enriched pathways (FDR < 0.1) were highlighted.

9-day fentanyl exposure in STBs produced transcriptional changes involving mitochondrial function and chromatin organization (Fig. 3D). These included decreased expression of *NDUFA4*, a vital component of the mitochondrial complex IV^55^, and multiple histone genes, including *H2BC7*, *H3C1*, *H3C2*, and *H3C11* (Table. S11). Notably, we found that the three genes related to trophectoderm activation and differentiation were affected by 9-day exposure, including *TRO* (log_2_FC: -6.02), which is implicated in trophoblast–endometrial adhesion^56^; *HAND1* (log_2_FC: 2.37), a transcription factor essential for trophoblast lineage specification and differentiation^57^; *IFITM3* (log_2_FC: -0.73), an inhibitor of villous trophoblast fusion^58^.

Pathway analysis demonstrated distinct transcriptional pattern in STBs exposed to fentanyl (Fig. 3E, Table S12, S13). Specifically, pathways related to mitochondrial function and ribosomal activity, including oxidative phosphorylation and translation-related processes, were consistently downregulated following 4-day or 9-day fentanyl exposure (right panel in Fig. 3E). In contrast, regulation of small GTPase-mediated signaling represented the only pathway commonly upregulated (Fig. 3E). Notably, none of these pathways were significantly enriched in TS-CTBs exposed to fentanyl (left panel in Fig. 3E). These results identified suppression of mitochondrial and protein-synthesis transcriptional programs as a prominent response to fentanyl exposure in STBs. Additionally, 4-day fentanyl exposure was linked to upregulation of cell junction assembly and cell cycle mitotic transition (Fig. 3E, Table S12), while 9-day exposure was characterized by downregulation of nucleosome-and chromatin-related pathways and upregulation of growth factor response pathways (Fig. 3E, Table S13).

### FMX co-exposure induces time-dependent transcriptional reprogramming characterized by early amplification and late divergence of fentanyl responses in STBs

We examined whether FMX co-exposure modified fentanyl-induced transcriptional responses in TS-STBs (Fig. 4A). Compared with 4-day exposure, 9-day FMX exposure led to a greater number of DEGs (Fig. 4B, Table S14, S15). Under 4-day FMX exposure, several genes associated with cellular stress responses were differentially expressed in STBs (Fig. 4C), including *DRAM1* (log_2_FC: -0.47), a mediator of DNA damage-induced autophagy, as well as the DNA damage response genes such as *PARP1* (log_2_FC: 0.44) and *MDC1* (log_2_FC: 0.62). We also found increased expression of *HAVCR1* (log_2_FC: 0.79), a receptor linked to trophoblast fusion^59^. Interestingly, several STB-dominant transcripts were also altered, including *PSG9* (log_2_FC: -0.45), *ALPP* (log_2_FC: -1.03), as well as *SIGLEC6* (log_2_FC: 0.50). Elevated expression of *SIGLEC6* is linked to mitochondrial dysfunction observed in preeclampsia^60^.

**Figure 4.**
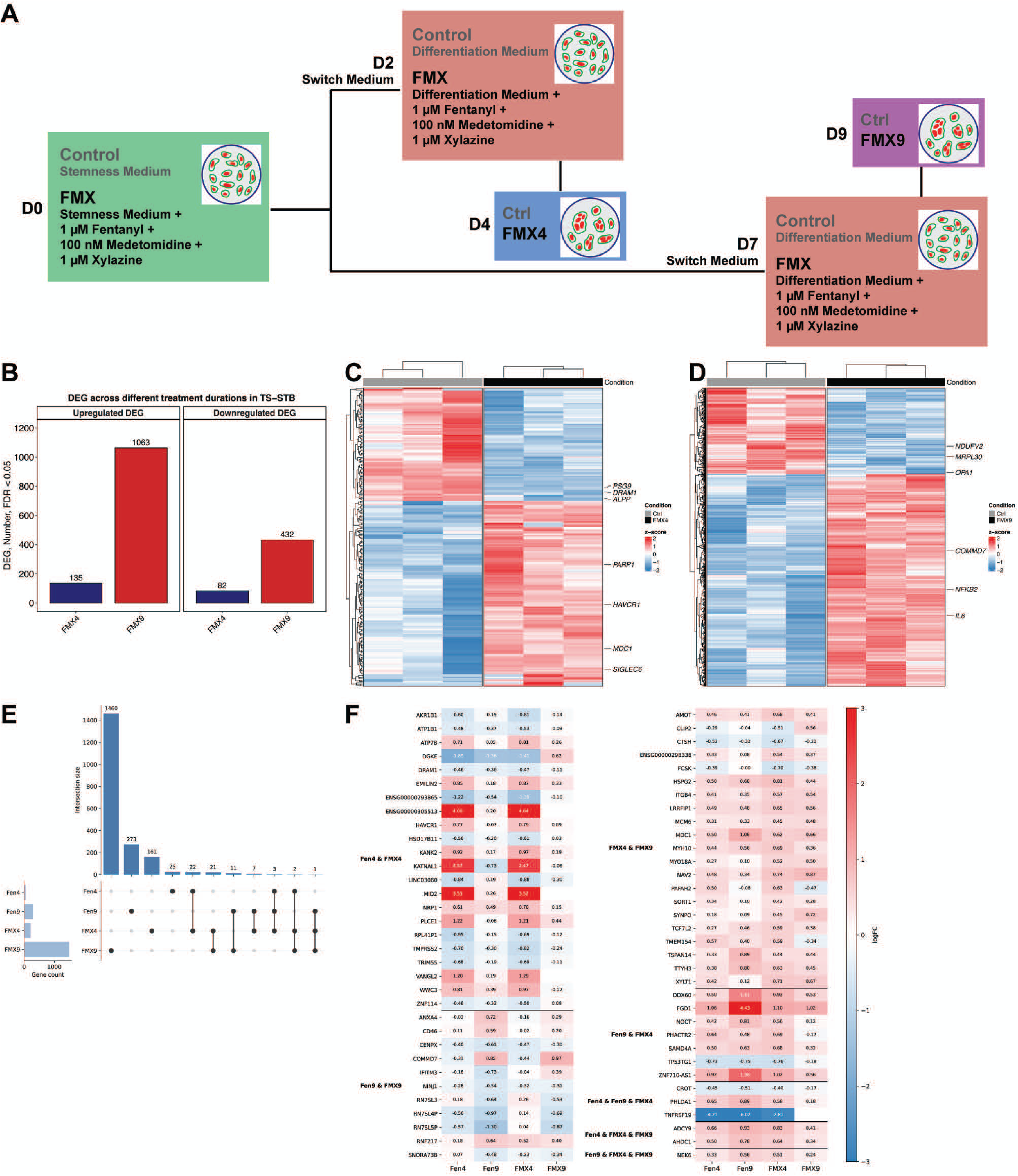
Effects of FMX co-exposure on transcriptional responses in STBs derived from TS-CTBs. (A) An experimental schematic: TS-CTBs were pre-exposed to 1 µM fentanyl, 100 nM medetomidine, and 1µM xylazine (FMX) in stemness medium (2 or 7 days), then transitioned to FMX-containing differentiation medium for 2 days to generate STBs representing 4-day (FMX4) and 9-day (FMX9) total exposure. (B) Barplot showing numbers of DEGs (FDR < 0.05) in STBs following 4-day and 9-day FMX exposure. (C–D) Heatmaps showing DEGs (FDR < 0.05) in STBs following (C) 4-day and (D) 9-day FMX exposure. (E) Upset plot showing overlapping DEGs between Fentanyl-and FMX-exposed STBs. (F) Heatmap showing the fold-change values of the overlapping DEGs identified in (E).

TS-STBs following 9-day FMX exposure led to gene expression changes, which were strongly associated with inflammatory responses and mitochondrial function (Fig. 4D). For instance, *COMMD7* (log_2_FC: 0.97), a regulator of NF-κB signaling pathway^61^, together with pro-inflammatory cytokine *IL6* (log_2_FC: 1.45) and *NFKB2* (log_2_FC: 0.57), an inducer of IL-6, were significantly upregulated, suggesting activation of pro-inflammatory pathways. Concurrently, genes that are critical for mitochondrial homeostasis were downregulated, including *NDUFV2*, *MRPL30*, and *OPA1*, suggesting suppression of mitochondrial transcriptional programs after extended FMX exposure.

We next compared overlapping DEGs across experimental groups. 4-day fentanyl and FMX shared 22 DEGs, whereas only 11 DEGs overlapped after 9-day exposure (Fig. 4E). All overlapping genes between Fen4 and FMX4 changed in the same direction, and most (16/22) exhibited a greater effect size in the FMX4 group (Fig. 4F, Fen4 & FMX4), including the stress-responsive genes *DRAM1* and *HAVCR1*. These findings suggest that FMX co-exposure amplifies early transcriptional responses to fentanyl. In contrast, among genes shared between Fen9 and FMX9, the magnitude of expression changes was generally attenuated in the FMX9 group (Fig. 4F, Fen9 & FMX9). Moreover, IFITM3 was downregulated following 9-day fentanyl exposure but upregulated following FMX exposure, suggesting that 9-day co-exposure does not merely intensify fentanyl-induced effects but instead reshapes the transcriptional program in STBs. Comparison of FMX4 and FMX9 uncovered 21 shared DEGs (Fig. 4E). Again, extending exposure duration did not uniformly increase transcriptional responses. Only four genes displayed progressive increase in expression levels, whereas most exhibited attenuated induction/suppression or even reversed direction over time (Fig. 4F, FMX4 & FMX9), highlighting the dynamic nature of adaptation to sustained FMX exposure.

GSEA pathway analysis further demonstrated that FMX co-exposure not only broadened the pathway-level alterations induced by fentanyl alone, but also shared a substantial proportion of transcriptional signatures (Fig. 5A). 4-day exposure (Fen4 or FMX4) resulted in enrichment of pathways related to mitotic cell cycle, histone modification, and hippo signaling, as well as downregulation of ribosomal RNA processing and metabolism, and ribosome biogenesis (Fig. 5B, Fen4 & FMX4). However, these patterns were largely reversed toward the baseline level following 9-day exposure. Pathways uniquely enriched following 4-day FMX exposure included those related to cell-cell junction organization, stress fiber assembly, nuclear DNA replication, and cytochrome complex, while 9-day FMX exposure was associated with enrichment of cadherin binding, RNA splicing, and peroxisome organization (Fig. 5B, FMX4 only and FMX9 only, Table S16, S17).

**Figure 5.**
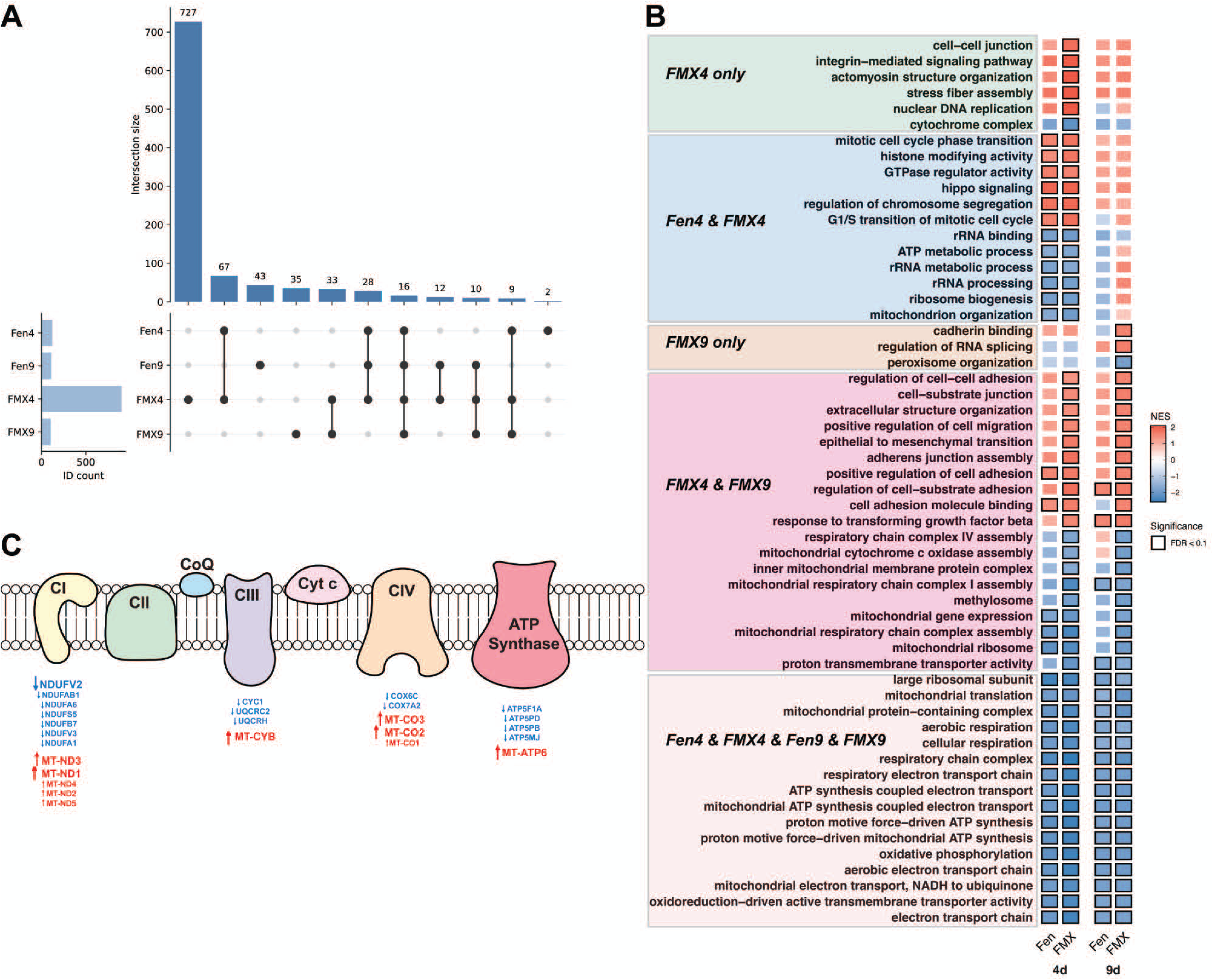
FMX co-exposure reprograms fentanyl-associated pathways in STBs derived from TS-CTBs. (A) Upset plot showing overlapping enriched pathways between Fentanyl-and FMX-exposed STBs. (B) Multi-group comparison of GSEA results in Fentanyl-and FMX-exposed STB cells at 4 days (left) and 9 days (right). The significantly enriched pathways (FDR < 0.1) were highlighted. (C) Expression of electron transport chain complex subunit genes. Genes differentially expressed in 9-day FMX-exposed STBs are annotated (FDR < 0.05).

FMX co-exposure at both time points enhanced pathways related to cell adhesion and junctions, epithelial mesenchymal transition, response to transforming growth factor, while strengthening suppression of transcriptional programs related to mitochondrial gene expression, mitochondrial translation, and respiratory chain complex assembly (Fig. 5B, FMX4 & FMX9).

A particularly notable finding in STBs was the consistent downregulation of mitochondrial transcriptional programs, including oxidative phosphorylation-related pathways, across all four exposure conditions (Fig. 5B, Fen4 & Fen9 & FMX4 & FMX9). Following 9-day FMX exposure, we found a pronounced imbalance between nuclear-encoded and mitochondrial-encoded genes that belong to the mitochondrial electron transport chain complexes. Specifically, nuclear-encoded mitochondrial genes were downregulated, whereas several mitochondrial genome-encoded transcripts were upregulated (Fig. 5C, Table S18).

### Fen and FMX induce cell-state-dependent transcriptional changes with limited gene-level convergence

We next compared the transcriptional profiles between drug-exposed TS-CTBs and derived STBs, and found limited overlap, observed only after 9-day fentanyl exposure (Fig. 6A). Two transcripts, *FLT1* and the lncRNA *ENSG00000269815,* were shared. Whereas *FLT1*, a hypoxia-responsive gene associated with oxidative stress and preeclampsia^62^, was elevated in both cellular states, *ENSG00000269815* showed opposite changes, being downregulated in STBs_Fen9 but upregulated in TS-CTBs_Fen9 (Fig. 6B). It is transcribed from *ABHD8* that is implicated in inhibiting innate immunity activation^63^.

**Figure 6.**
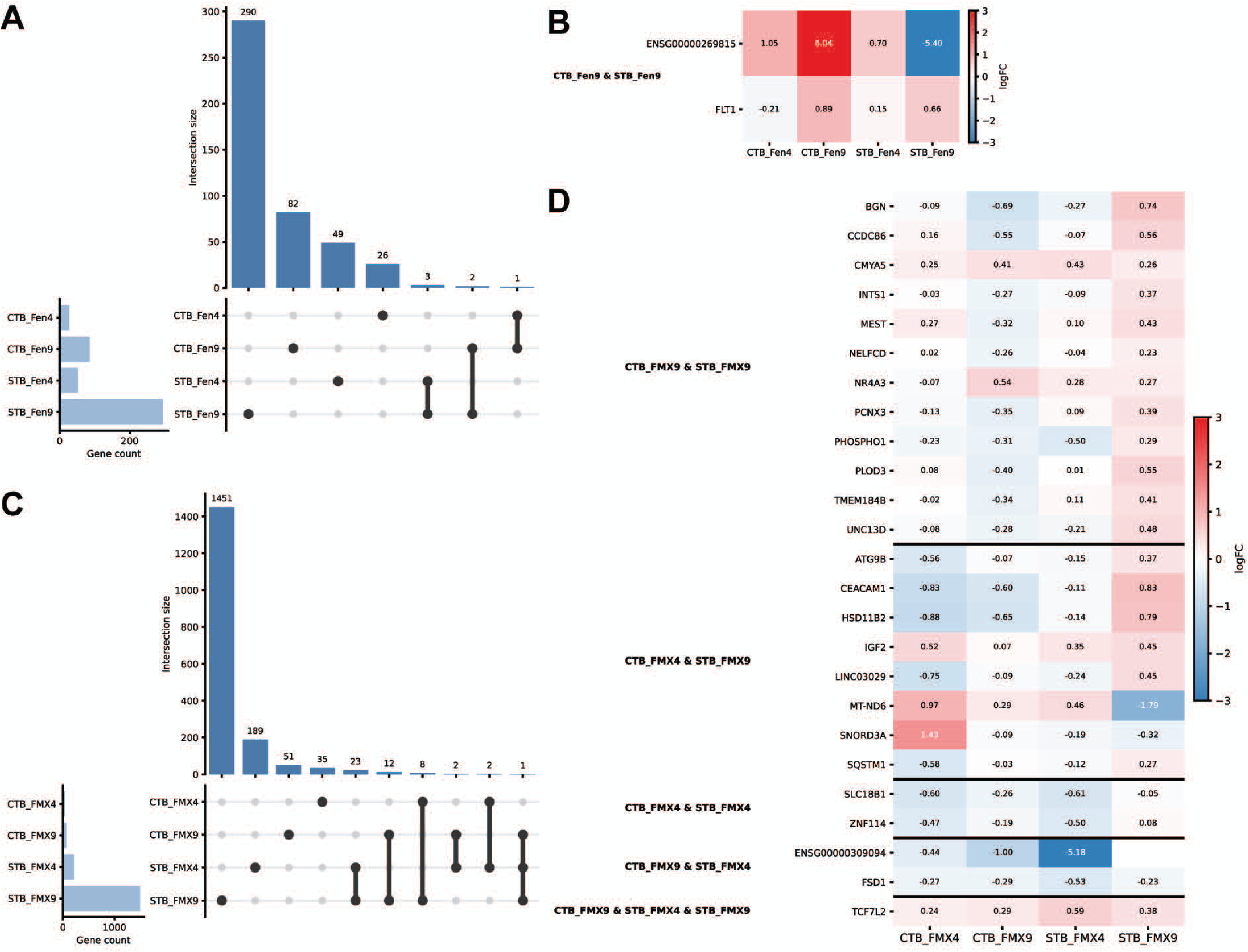
Comparison of transcriptional responses of TS-CTBs vs derived STBs following Fentanyl and FMX exposure. (A and C) Upset plot showing overlapping DEGs between TS-CTBs and STBs exposed to (A) fentanyl and (C) FMX. (B and D) Heatmaps showing fold-change values of the overlapping DEGs in (B) Fentanyl-and (D) FMX-exposed groups.

In contrast to fentanyl alone, FMX exposure resulted in a larger number of shared DEGs between TS-CTBs and derived STBs (Fig. 6C). However, the majority of overlapping DEGs showed opposite direction of expression changes in TS-CTBs and derived STBs (Fig. 6D), suggesting that some shared molecular targets exhibit cell-state-dependent transcriptional responses. Notably, transcription factor TCF7L2 is persistently upregulated in TS-CTBs and derived STBs exposed to the FMX (Fig. 6D, CTB_FMX9 & STB_FMX4 & STB_FMX9), whereas ZNF114 is downregulated in both TS-CTBs and STBs exposed to 4-day FMX (Fig. 6D, CTB_FMX4 & STB_FMX4). Together, these findings indicate that TS-CTBs and STBs exhibit fundamentally distinct transcriptional responses to fentanyl and FMX co-exposure.

### Pathway analysis shows exposure-duration-dependent transcriptional responses to fentanyl and FMX exposure in TS-CTBs and derived STBs

To further examine the dynamic effects of fentanyl alone or the three drug FMX combination on TS-CTBs and derived STBs, we performed gene set variation analysis (GSVA) to visualize the magnitude of changes in pathway activity, defined by GSVA scores. Positive GSVA scores indicated increased pathway enrichment relative to control, whereas negative scores indicated reduced enrichment relative to control. Unsupervised clustering revealed that transcriptional responses were primarily organized according to exposure type and duration, with pathways displaying distinct temporal patterns of activation or suppression (Fig. 7, Table S19).

**Figure 7.**
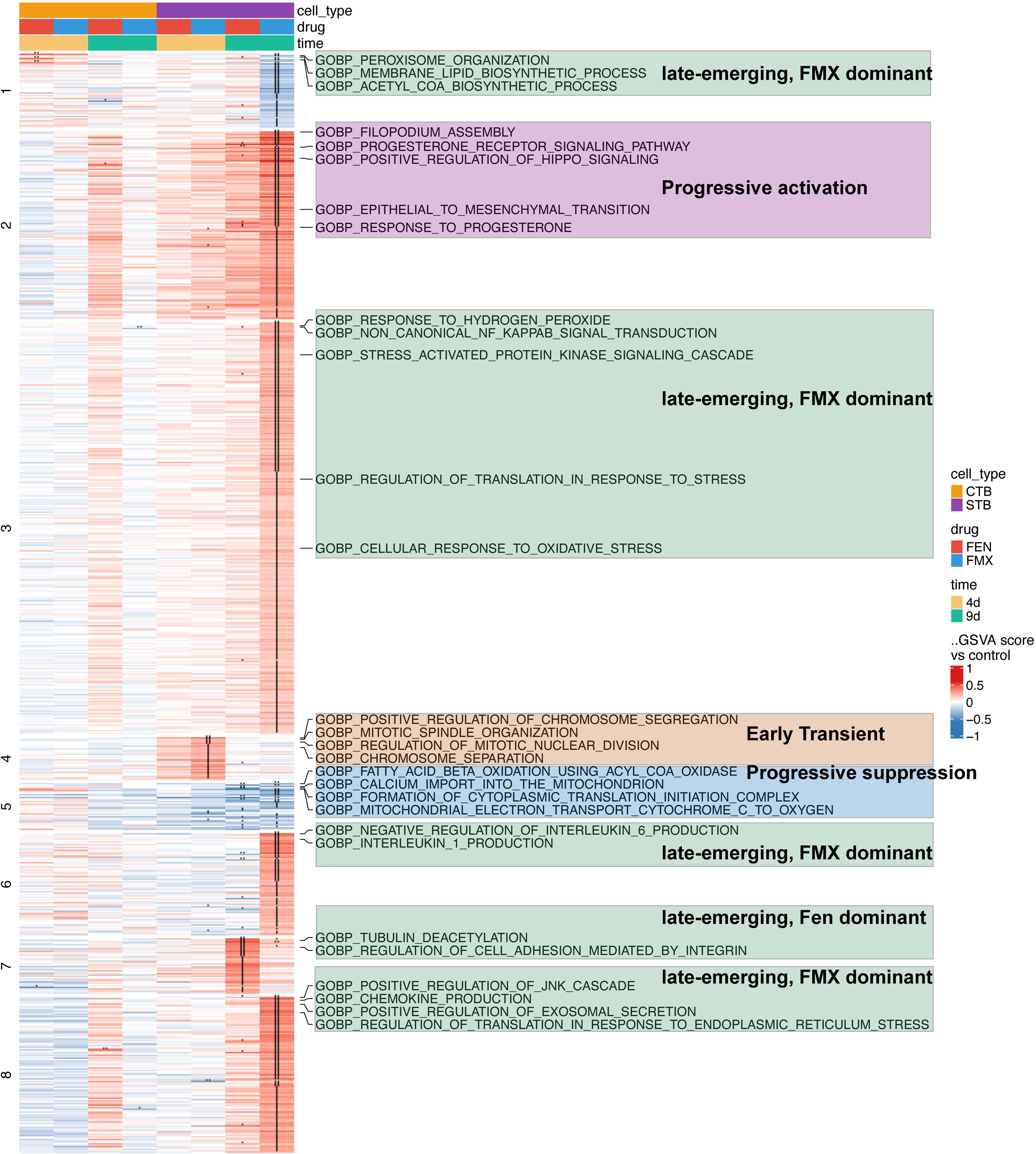
Distinct activity programs induced by drug exposure in trophoblasts. Heatmap showing changes in clustered pathway activity, inferred by GSVA (see text). Pathway activities were quantified at the sample level and compared among each drug-exposed group and its matched control within the same cell state and time point. Color scale represents the difference in GSVA enrichment score relative to control. Pathways shown were selected with adjusted p value < 0.1 in at least one contrast and clustered by activity patterns across all conditions. Representative pathways from each cluster are labeled on the right.

We first assessed pathways that exhibited progressive and consistent changes over time (Fig.7, cluster 2 and 5). Cluster 2 comprised pathways related to progesterone receptor signaling, response to progesterone, positive regulation of Hippo signaling, filopodium assembly, and epithelial-to-mesenchymal transition. These pathways initially showed modestly positive enrichment with 4-day exposure, with stronger activation after 9-day exposure. Unlike in Cluster 2, pathways in Cluster 5 contained those associated with mitochondrial function and cellular oxidation that showed early transcriptional suppression, which became more pronounced after 9 day of drug exposure. These findings highlight that 9-day drug exposure is characterized by simultaneous activation of trophoblast remodeling pathways and progressive repression of mitochondrial-related transcriptional programs.

We next examined pathways that were transiently altered following 4-day exposure but returned toward baseline after 9-day exposure (Fig. 7, Cluster 4). These pathways were enriched for mitotic cell cycle processes, specifically chromosome segregation and separation, indicating an early and transient effect of drug exposure on trophoblast proliferative programs.

Finally, we focused on pathways that emerged only after 9-day exposure. In STBs exposed to fentanyl alone, late-response pathways were primarily associated with cell adhesion and tubulin deacetylation (Fig. 7, Cluster 7), suggesting that 9-day fentanyl exposure drives cytoskeletal remodeling and adhesion pathway activation. In contrast, 9-day FMX co-exposure selectively suppressed pathways related to lipid biosynthesis and peroxisome organization (Fig. 7, Cluster 1). Additionally, pathways related to cellular stress, signaling pathways, immune regulation, and exosomal secretion (Fig. 7, Cluster 3, 6, 8) were only enriched in STBs with 9-day FMX co-exposure, suggesting that broader engagement of adaptive and remodeling programs following combined drug exposure.

## DISCUSSION

We interrogated the transcriptomic landscape of human trophoblasts following fentanyl exposure alone or in combination with xylazine and medetomidine, two major adulterants detected in the illicit fentanyl market. Using the well-defined human TS-CTBs and derived STB model system, we illustrate a cell-type resolved overview of their responses to drug exposure. Three major findings are supported by our data: First, trophoblastic cell states exhibit distinct transcriptional changes to fentanyl exposure, with stronger perturbations in STBs than parental TS-CTBs. Second, both fentanyl and FMX exposure induce suppression of mitochondrial oxidative phosphorylation-related transcriptional programs while promoting progressive trophoblast remodeling programs in TS-CTB-derived STBs. Third, co-exposure to xylazine and medetomidine reshapes fentanyl-induced transcriptomic alteration in a time-specific manner.

Our data highlight the greater transcriptional sensitivity of STBs compared to TS-CTBs. 4-day fentanyl exposure induced relatively limited changes in TS-CTBs, whereas derived STBs displayed broader alterations affecting metabolic and translational pathways. This difference likely reflects fundamental biological distinctions of the two cellular states. Differentiation from TS-CTBs to STBs involves extensive metabolic remodeling and increased biosynthetic demand that is essential for hormone production and maternal-to-fetal nutrition supply. These biological activities fulfilled by STBs may underline their susceptibility to disruptions in stress-response and metabolic pathways, induced by drug exposure.

One of the most consistent transcriptional signatures was suppression of mitochondrial transcriptional programs, particularly gene expression associated with mitochondrial electron transport chain (ETC) complexes, in STBs exposed to fentanyl alone or to the FMX mixture. Mitochondrial function plays a central role in placental energy metabolism, and its dysfunction has been implicated in major pregnancy complications^64,65^. Combined FMX exposure led to additional transcriptional disturbance, manifested by discordant expression control of nuclear-encoded and mitochondrial genome-encoded genes of the ETC complexes. This mito-nuclear imbalance^66^ defines loss of coordinated transcriptional regulation between the nuclear and mitochondrial genomes and may comprise mitochondrial homeostasis.

Our GSVA analysis indicates that fentanyl induces time-dependent transcriptional changes in trophoblasts, with early pathway shifts at day 4 preceding additional remodeling at day 9. Early activation of hormone-responsive pathways may reflect a priming state that defines subsequent cellular adaptation. With extended exposure, pathway activity shifts toward cytoskeletal, adhesion, and secretion-related programs, accompanied by progressive inhibition of metabolic and mitochondrial transcriptional programs, a pattern that aligns with altered cellular metabolic homeostasis. Nevertheless, FMX co-exposure does not uniformly amplify fentanyl effects, but instead reprogrammes pathway responses in a cell type-dependent manner.

Our results suggest that combined exposure elicits a broader transcriptional response than fentanyl alone, highlighting potential crosstalk between opioid and ɑ2-adrenergic signaling pathways. Supporting this possibility, xylazine has been reported to act as an agonist of kappa opioid receptor^67^, and to enhance fentanyl-associated lethality in mice^68^. Together, these findings indicate that opioid and adrenergic pathways may interact at multiple levels, with underlying mechanisms warranting further investigation.

Limitations should be considered. First, the findings do not capture donor-to-donor variability or fully recapitulate the cellular complexity of the human placenta. Validation in additional TS cell lines and primary cells will therefore be required to define the generalizability of the identified transcriptional responses. Exposure was limited to a 2-day differentiation window, chosen to assess transcriptional responses during differentiation. The effects of drug exposure across the full course of STB differentiation will require future study. Single-nucleus analysis of placental biopsies could further resolve transcriptional programs underlying real-world opioid polysubstance exposure. Second, differences in receptor expression between TS-CTBs and STBs have not been fully characterized, and the contribution of receptor abundance and signaling to cell-state-specific transcriptional responses remains unclear. Third, the transcriptomic changes identified herein point to functional processes for further investigation, including mitochondrial function and trophoblast remodeling. Replicate-level analyses assess consistency and variability within the current RNA-seq dataset, but do not constitute validation in an independent cohort. Differences between the two exposure durations should not be interpreted as validated temporal trajectories. Studies using additional biological samples, qRT-PCR, protein-level assays, and targeted functional experiments will be necessary to establish the reproducibility and functional significance of selected transcriptomic changes. Fourth, because individual adulterant-only exposure groups were not included, the responses observed under FMX co-exposure cannot be independently attributed to either adulterant, nor can potential additive, antagonistic, or synergistic interactions be inferred. Accordingly, the findings are interpreted as responses to the complete FMX mixture. Future studies that incoporate individual adulterant and factorial co-exposure conditions will be necessary to distinguish the contributions and interactions of these agents. Finally, although drug concentrations were informed by available human toxicology reports^18–20^, real-world exposure patterns are highly variable among individuals and continue to evolve with changes in illicit drug markets.

Integrating dose-response and time-resolved experiments with placental exposure data will be important for evaluating the translational relevance of these findings. This study provides a discovery framework for prioritizing subsequent functional validation and for examining how polysubstance exposure alters trophoblast metabolism and endocrine function.

## Availability of data and materials

RNAseq data were deposited in NCBI Sequence Read Archive with BioProject IDs PRJNA1506347, PRJNA1506349, and PRJNA1506350.

## Author contributions

Y.O. and Y.S. conceived the project; Y.O. and Y.S. designed the study; Y.O., J.M., L.L., K.H. performed the experiments; Y.O., S.L., J.M. analyzed the data; Y.O., J.M., S.L., and Y.S. reviewed the data and interpreted the results. J.M., Y.O. and Y.S. wrote the manuscript. All the authors reviewed the manuscript.

## Acknowledgements

We thank Takahiro Arima and the RIKEN Cell Bank for providing us with TS cells.

## Abbreviations

*ABHD8*: Abhydrolase domain containing 8
*ADAM*: A disintegrin and metalloprotease
*ALPP*: Alkaline phosphatase, placental
*CEACAM1*: CEA cell adhesion molecule 1
*CHRNA3*: Cholinergic receptor nicotinic alpha 3 subunit
*CGB3*: Chorionic gonadotropin subunit beta 3
*CGB5*: Chorionic gonadotropin subunit beta 5
*CGB7*: Chorionic gonadotropin subunit beta 7
*CGB8*: Chorionic gonadotropin subunit beta 8
*COLEC12*: Collectin subfamily member 12
*COMMD7*: COMM domain containing 7
*CROT*: Carnitine O-octanoyltransferase
DEG: Differentially expressed genes
*DRAM1*: DNA damage regulated autophagy modulator 1
EMT: Epithelial mesenchymal transition
Fen: Fentanyl
Fen4: Fentanyl exposure for 4 days
Fen9: Fentanyl exposure for 9 days
*FLT1*: Fms related receptor tyrosine kinase 1
FMX: Fentanyl in combination with medetomidine and xylazine
FMX4: Fentanyl in combination with medetomidine and xylazine for 4 days
FMX9: Fentanyl in combination with medetomidine and xylazine for 9 days
*GCM1*: Glial cells missing 1
*H2BC7*: H2B clustered histone 7
*H3C1*: H3 clustered histone 1
*H3C11*: H3 clustered histone 11
*H3C2*: H3 clustered histone 2
*HAND1*: Heart and neural crest derivatives expressed 1
*HAVCR1*: Hepatitis A virus cellular receptor 1
*HSD11B2*: Hydroxysteroid 11-beta dehydrogenase 2
*HSD17B11*: Hydroxysteroid 17-beta dehydrogenase 11
*IFITM3*: Interferon induced transmembrane protein 3
*IGF2*: Insulin like growth factor 2
*IL6*: Interleukin 6
*lncRNA*: Long non-coding RNA
*LGALS13*: Lectin, Galactoside-Binding, Soluble, 13
*MDC1*: Mediator of DNA damage checkpoint 1
*MRPL30*: Mitochondrial ribosomal protein L30
*NCF1*: Neutrophil cytosolic factor 1
*NDUFA4*: NADH:ubiquinone oxidoreductase subunit A4
*NDUFAF1*: NADH:ubiquinone oxidoreductase complex assembly factor 1
*NDUFV2*: NADH:ubiquinone oxidoreductase core subunit V2
*NFKB2*: Nuclear factor kappa B subunit 2
*NR1H4*: Nuclear receptor subfamily 1 group H member 4
*NR4A3*: Nuclear receptor subfamily 4 group A member 3
*OPA1*: OPA1 mitochondrial dynamin like GTPase
*OVOL1*: Ovo like transcriptional repressor 1
*PARP1*: Poly (ADP-ribose) polymerase 1
*PHLDA1*: Pleckstrin homology like domain family A member 1
*PRDM1*: PR/SET domain 1
*PSG3*: Pregnancy specific beta-1-glycoprotein 3
*PSG9*: Pregnancy specific beta-1-glycoprotein 9
*SGK1*: Serum/glucocorticoid regulated kinase 1
*SIGLEC6*: Sialic acid binding Ig like lectin 6
*SQSTM1*: Sequestosome 1
*SRD5A3*: Steroid 5 alpha-reductase 3
STB: Syncytiotrophoblast
*SYDE1*: Synapse defective protein 1
*TCF7L2*: Transcription factor 7 like 2
*TNFRSF19*: TNF receptor superfamily member 19
*TRO*: Trophinin
TS: Trophoblast stem

