## Supplementary figures for "Fentanyl and fentanyl-xylazine co-exposure influence mitochondrial transcriptional programs in human trophoblasts"

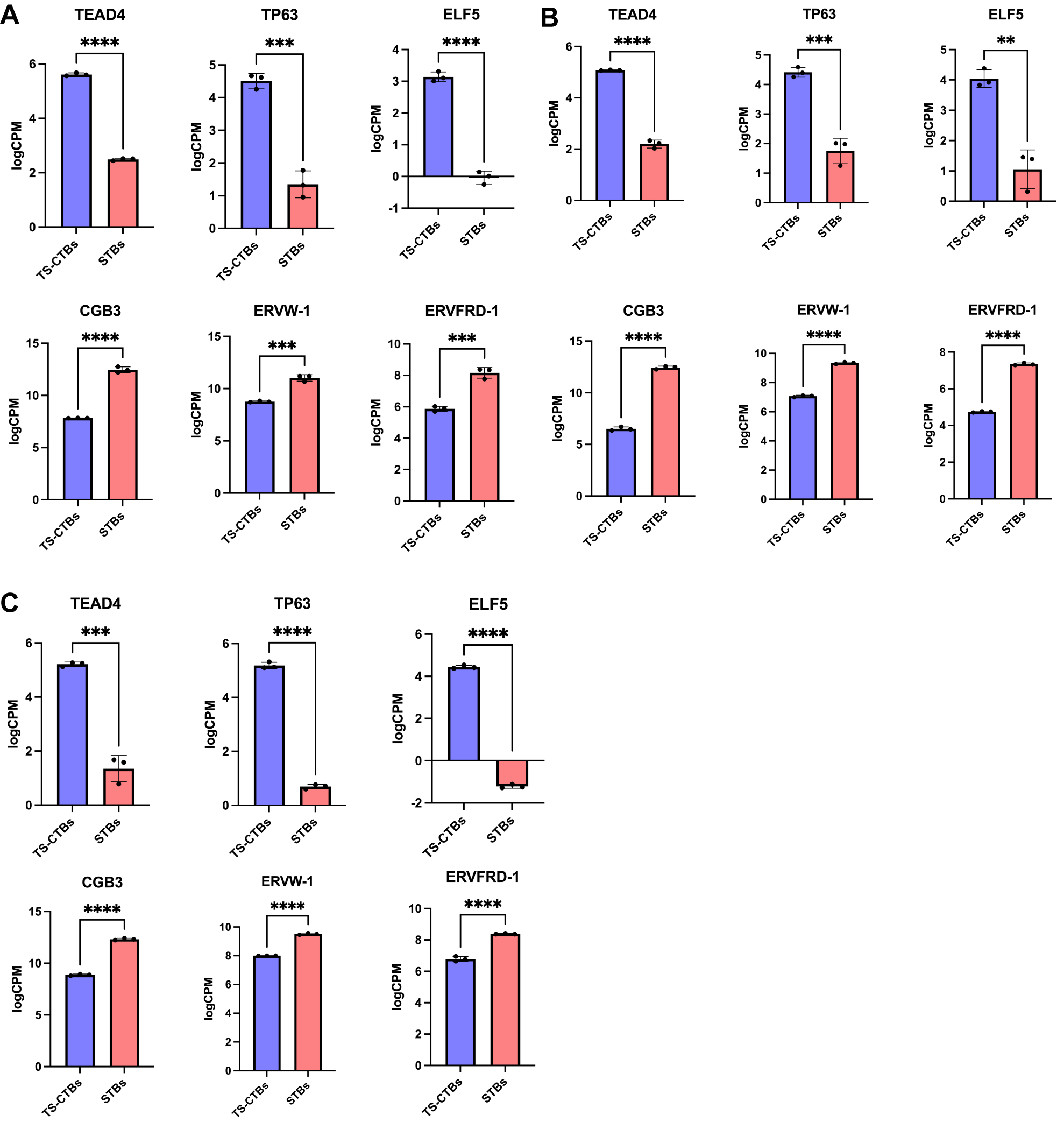

***Figure S1. The transcript levels of TS-associated and STB-associated genes in TS-CTBs and derived STBs.*** TS-associated genes include *TEAD4*, *TP63*, and *ELF5*, whereas STB-associated genes include *CGB3*, *ERVW-1*and *ERVFRD-1*. (A) Gene expression in vehicle-control TS-CTBs and derived STBs at day 4. (B) Gene expression in vehicle control samples from the 9-day Fen experiment. (C) Gene expression in vehicle control samples from the 9-day FMX experiment. FDR < 0.05, n=3 biological replicates.

Fig. S1: We selected 2-day differentiation window to match the period in which syncytial fusion is initiated, and cells begin acquiring endocrine function. We did not extend it to 5–6 days because mature STBs show prominent senescence-associated features^1^, and syncytial degeneration has been reported in cultured villi^2^. These factors may confound drug effects. The effects of prolonged exposure across the full 5–6-day differentiation were not examined and will require future study, which is acknowledged as one of the limitations of this study.

In our system, 2 days of differentiation was sufficient to initiate the transition from TS-CTB to STB. Canonical trophoblast stemness genes (ELF5, TEAD4, TP63) were downregulated, and STB-associated genes (CGB3, ERVW-1, ERVFRD-1) were upregulated (Fig. S1), supporting that this window captures early STB differentiation. Expression is shown as Trimmed Mean of M values (TMM)-normalized log2 counts per million (logCPM); negative values correspond to CPM < 1 and reflect low transcript abundance. This differentiation protocol has also been validated in our previous studies^3^.

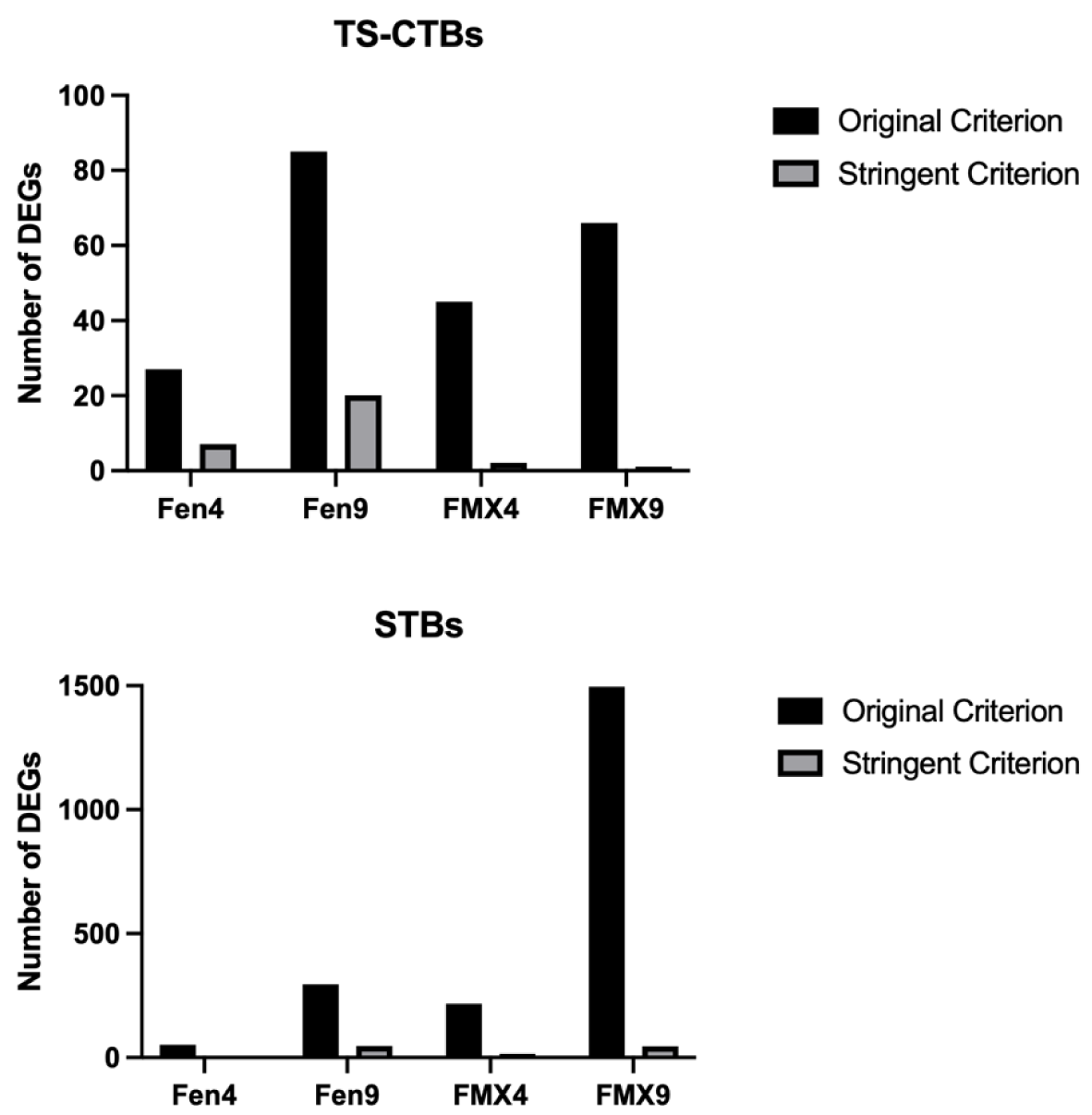

***Figure S2. Comparison of DEG counts identified using the original and stringent significance criteria. The original criterion was FDR < 0.05, and the stringent criterion was FDR < 0.01 with an absolute fold change ≥ 2.*** (A) Number of DEGs in TS-CTBs after 4 or 9 days of exposure to Fen or FMX. (B) Number of DEGs in STBs after 4 or 9 days of exposure to Fen or FMX.

Fig. S2: We used FDR < 0.05 as the criterion for the differentially expressed genes (DEGs) analysis. This threshold is a commonly used cutoff in RNA-seq differential-expression analyses^4^. We agree that a more stringent threshold is useful for assessing the robustness of the DEG findings. We therefore performed an additional sensitivity analysis using FDR < 0.01 and an absolute fold change ≥ 2. This substantially reduced the number of DEGs (Fig. S2), particularly in the TS-CTB comparisons, and most of the gene-level overlaps identified using the original criteria were no longer retained. Given the small number of DEGs under this threshold, DEG-based GO enrichment analysis was not informative. Based on these results, we retained FDR < 0.05 as the primary criterion for DEG analysis. Moreover, GSEA and GSVA were used as complementary pathway-level approaches to evaluate coordinated transcriptional changes.

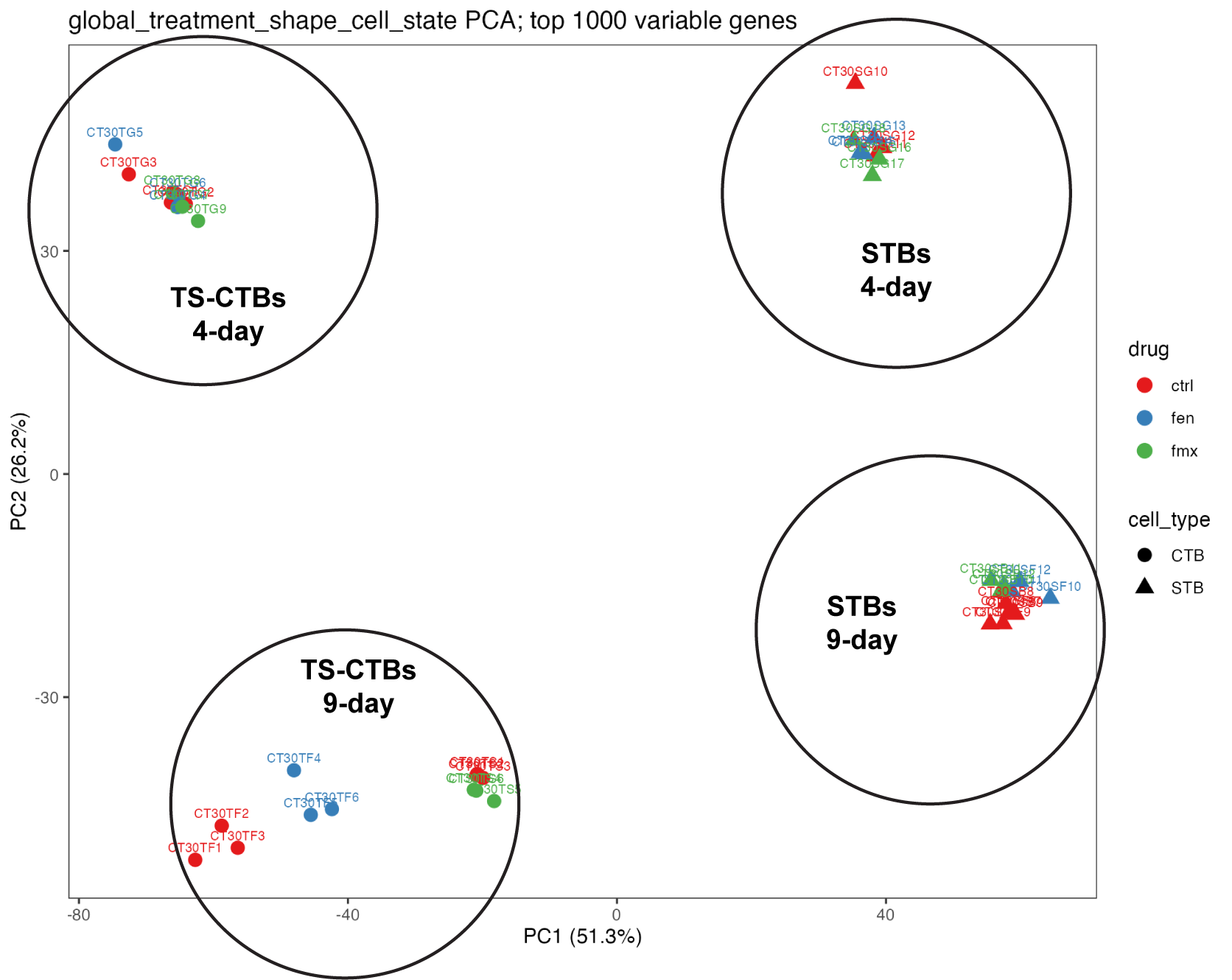

***Figure S3. PCA of all bulk RNAseq samples.*** PCA was performed on variance-stabilized expression values for the 1,000 most variable genes across all 42 samples. Samples separate primarily by cell state and exposure duration. Points are shaped by cell state and colored by drug exposure. PC1 and PC2 explain 51.3% and 26.2% of the variance, respectively.

Fig. S3: a sample-level analysis provides an overview of drug effects. The RNA-seq dataset comprises three independent experimental batches. Each experimental conditions included three biological replicates, defined as independently cultured, drug-exposed samples, with matched vehicle controls included in every batch.

The RNA-seq datasets comprise 42 samples from three independent experimental batches, each with its own vehicle controls and drug-exposed biological replicates (n=3 per condition).

Batch 1 included TS-CTBs and STBs exposed for 4 days to vehicle, fentanyl (Fen), or the fentanyl/medetomidine/xylazine combination (FMX).

Batch 2 included TS-CTBs and STBs exposed for 9 days to vehicle or Fen.

Batch 3 included TS-CTBs and STBs exposed for 9 days to vehicle or FMX.

Vehicle controls were matched to each cell state and exposure duration within each batch.

To assess consistency and the overall relationships across all 42 samples, we performed pairwise Pearson correlation analysis and global principal component analysis (PCA) using genes with >10 read counts in at least three samples. Pearson correlation coefficients among replicates within the same condition ranged from 0.92 to 0.99, indicating high concordance of the transcriptomic profiles across replicates (Table S20). In the global PCA, replicated from the same condition generally clustered together, while sample distribution reflected the combined influence of trophoblast cell state, exposure duration, drug type, and experimental batch (Fig. S3). Because the global PCA encompassed multiple biological and experimental factors, we used it as an overview of dataset structure rather than as evidence of drug-specific effects. Drug-associated differential gene expression and pathway analyses were therefore conducted within matched comparisons. No samples were excluded.

**Replicate-level analyses** include a pairwise sample-correlation heatmap and a clustered heatmap of differentially expressed genes. These analyses allow direct assessment of agreement and variability among the biological replicates included in the RNA-seq datasets.

A pairwise sample-correlation heatmap across all samples was first generated to assess agreement among biological replicates and to visualize relationships among samples across trophoblast cell states, treatment groups, and exposure durations (Fig. S4).

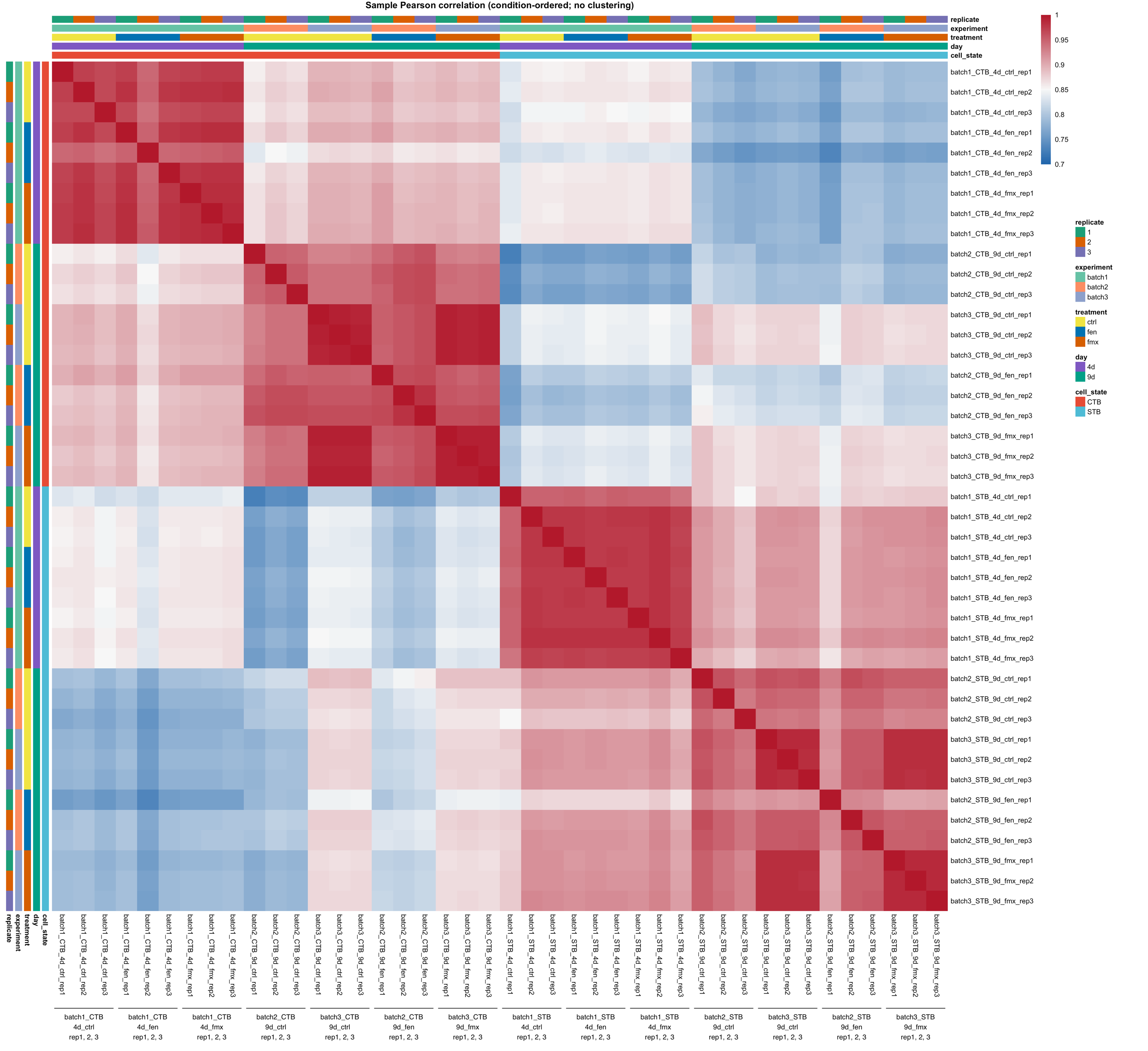

***Figure S4****.* ***Pairwise correlations among RNA-seq samples.*** *Heatmap showing pairwise Pearson correlation coefficients across all RNA-seq samples. Samples are annotated by trophoblast cell state, treatment condition, and exposure duration.*

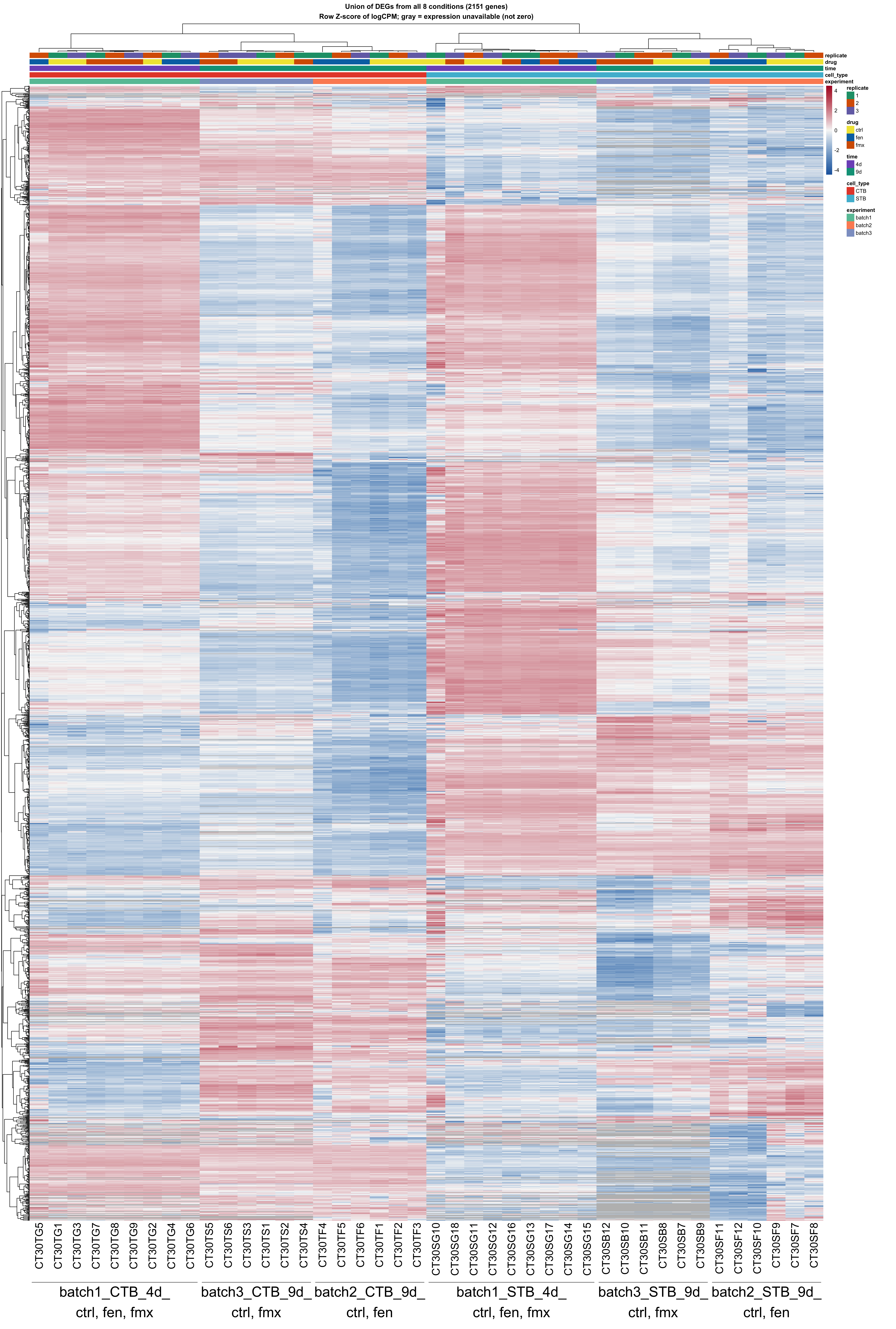

***Figure S5. Expression patterns of significantly differentially expressed genes across biological replicates and experimental conditions.*** *Hierarchical clustered heatmap showing the expression of genes identified as differentially expressed at FDR < 0.05 in at least one prespecified treatment-versus-matched-control comparison at day 4 or day 9. Columns represent individual biological replicates and are annotated by trophoblast cell state, treatment group, and exposure duration. Rows represent differentially expressed genes.*

A hierarchically clustered heatmap showing the expression of significantly differentially expressed genes (DEG) across individual biological replicates and experimental conditions. The heatmap enables visualization of both shared and variable DEG expression patterns among individual replicates. Sample organization was largely associated with trophoblast cell state and exposure duration, while the degree of treatment-associated separation varied among comparisons. We therefore evaluated differential gene expression and pathway enrichment results *within each prespecified treatment-versus-matched-control comparison* rather than inferring treatment effects from global clustering alone.

Together, these additional analyses (Fig. S4, S5) provide a transparent assessment of replicate-level consistency and variability in the RNA-seq dataset.

**Supplementary notes**

The primary aim of this study was to profile transcriptome-wide responses to fentanyl and FMX across trophoblast states and exposure durations, rather than to validate a limited set of candidate genes, we therefore focused our interpretation on transcriptome-wide and pathway-level findings. qRT-PCR using the same samples would provide technical confirmation but would not demonstrate reproducibility in an independent biological cohort. Because no independent validation cohort was included in the initial experimental design, we did not add qRT-PCR validation of selected genes. RNA-seq is itself a well-established and reliable method for measuring transcript abundance and differential expression, and benchmarking studies have shown good concordance between RNA-seq and qRT-PCR results^5,6^ .

We interpreted the DEG results together with pathway-level analyses (GSEA and GSVA), which assess coordinated transcriptional changes rather than individual genes, and we did not attribute physiological significance to individual DEGs. Our transcriptomic analyses *showed no significant enrichment* of gene sets related to cell morphology, proliferation, or apoptosis (FDR > 0.1). These data therefore do not indicate overt changes in these processes. However, transcript-level data cannot exclude phenotypic effects. The objective of this study was to define transcriptional responses to fentanyl and FMX across trophoblast culture states and exposure durations. The observed pathway-level changes provide a transcriptomic framework for identifying candidate processes for future functional investigation.

The objective of this study was to characterize drug exposure-associated transcriptional changes in human trophoblast cells exposed to fentanyl alone or in combination with medetomidine and xylazine, rather than to establish phenotypic impairment or causal mechanisms. The RNA-seq findings identify changes in gene expression and biological pathways that generate testable hypotheses regarding cellular responses to these exposures. However, transcriptomic changes do not establish corresponding alterations in protein abundance, cellular function, or physiological outcomes. Validation using additional biological samples and orthogonal assays is acknowledged as an important direction for future investigation.

1	Gal, H., Lysenko, M., Stroganov, S. *et al.* Molecular pathways of senescence regulate placental structure and function. *EMBO J* **38**, e100849, doi:10.15252/embj.2018100849 (2019).

2	Palmer, M. E., Watson, A. L. & Burton, G. J. Morphological analysis of degeneration and regeneration of syncytiotrophoblast in first trimester placental villi during organ culture. *Hum Reprod* **12**, 379-382, doi:10.1093/humrep/12.2.379 (1997).

3	Sadovsky, E., Chu, T., Barak, O. *et al.* The impact of opioids on the transcriptional landscape of human villous trophoblasts. *Placenta* **143**, 54-61, doi:10.1016/j.placenta.2023.10.001 (2023).

4	McCarthy, D. J., Chen, Y. & Smyth, G. K. Differential expression analysis of multifactor RNA-Seq experiments with respect to biological variation. *Nucleic Acids Res* **40**, 4288-4297, doi:10.1093/nar/gks042 (2012).

5	Coenye, T. Do results obtained with RNA-sequencing require independent verification? *Biofilm* **3**, 100043, doi:10.1016/j.bioflm.2021.100043 (2021).

6	Consortium, S. M.-I. A comprehensive assessment of RNA-seq accuracy, reproducibility and information content by the Sequencing Quality Control Consortium. *Nat Biotechnol* **32**, 903-914, doi:10.1038/nbt.2957 (2014).
